## Supplemental Material for "rRNA methylation by Spb1 regulates the GTPase activity of Nog2 during 60S ribosomal subunit assembly"

### Supplemental Table S1 - Cryo-EM data collection, refinement and validation statistics

| | Nog2$^{pre}$ *SPB1* (emdb-26651) (pdb 7UOO) | Nog2$^{pre}$ 5S rRNP *SPB1* (emdb-26689) | Nog2$^{pre}$ s*pb1*$^{D52A}$ (emdb-26703) (pdb 7UQZ) | Nog2$^{post}$ *spb1*$^{D52A}$ (emdb-26799) (pdb 7UUI) | Nog2$^{pre}$-AlF$_4^-$ *spb1*$^{D52A}$ (emdb-26686) (pdb 7UQB) | Nog2$^{pre}$ *spb1*$^{D52A/E769E}$ (emdb-26941) (pdb 7V08) |
|---|---|---|---|---|---|---|
| **Data collection and processing** | | | | | | |
| Magnification | 81,000x | 81,000x | 81,000x | 81,000x | 81,000x | 81,000x |
| Voltage (kV) | 300 | 300 | 300 | 300 | 300 | 300 |
| Electron exposure (e$^-$/Å$^2$) | 56 | 56 | 56 | 56 | 50 | 56 |
| Defocus range (μm) | -0.9 – (-2.2) | -0.9 – (-2.2) | -0.9 – (-2.2) | -0.9 – (-2.2) | -0.9 – (-2.2) | -0.9 – (-2.2) |
| Pixel size (Å) | 1.08 | 1.08 | 1.08 | 1.08 | 1.07 | 1.08 |
| Symmetry imposed | C1 | C1 | C1 | C1 | C1 | C1 |
| Initial particle images (no.) | 1,639,317 | 1,639,317 | 905,023 | 905,023 | 716,018 | 1,207,583 |
| Final particle images (no.) | 1,159,206 | 1,035,794 | 328,470 | 50,091 | 417,994 | 320,145 |
| Map resolution (Å) | 2.34 | 2.63 | 2.44 | 2.90 | 2.38 | 2.36 |
| FSC threshold | 0.143 | 0.143 | 0.143 | 0.143 | 0.143 | 0.143 |
| Map resolution range (Å) | 2.2 – 10 | 2.2 – 10 | 2.2 – 10 | 2.2 – 10 | 2.2 – 10 | 2.2 – 10 |
| **Refinement** | | | | | | |
| Initial model used (PDB code) | 3JCT | | 3JCT | 6YLH | 3JCT | 3JCT |
| Model resolution (Å) | 2.52 | | 2.60 | 3.33 | 2.53 | 2.48 |
| FSC threshold | 0.5 | | 0.5 | 0.5 | 0.5 | 0.5 |
| Map sharpening *B* factor (Å$^2$) | 79.52 | | 82.93 | 87.17 | 70.61 | 61.93 |
| Model composition | | | | | | |
| Non-hydrogen atoms | 157949 | | 157839 | 3672 | 157949 | 157849 |
| Protein residues | | | | | | |
| Nucleotides | 10806 | | 10806 | 452 | 10806 | 10806 |
| | 3329 | | 3325 | 1 | 3330 | 3325 |
| *B* factors (Å$^2$) | | | | | | |
| Protein | 22.13 | | 19.42 | 21.70 | 19.17 | 21.39 |
| Nucleotides | 23.34 | | 21.14 | 17.29 | 20.90 | 32.00 |
| Ligand | 15.98 | | 11.19 | 12.13 | 12.04 | 12.91 |
| Waters | 16.23 | | 11.01 | | 10.89 | 8.39 |
| R.m.s. deviations | | | | | | |
| Bond lengths (Å) | 0.002 | | 0.002 | 0.002 | 0.003 | 0.005 |
| Bond angles (°) | 0.563 | | 0.536 | 0.504 | 0.576 | 0.729 |
| Validation | | | | | | |
| MolProbity score | 1.31 | | 1.23 | 1.13 | 1.28 | 1.29 |
| Clashscore | 3.41 | | 3.67 | 3.38 | 4.24 | 4.08 |
| Poor rotamers (%) | 0.2 | | 0.33 | 0 | 0.43 | 0.55 |
| Ramachandran plot | | | | | | |
| Favored (%) | 97.56 | | 97.65 | 98.65 | 97.68 | 97.50 |
| Allowed (%) | 2.44 | | 2.35 | 1.35 | 2.32 | 2.50 |
| Disallowed (%) | 0 | | 0 | 0 | 0 | 0 |

**Supplemental Table S2 - Yields of Nog2·Tif6 pre-60S ribosomes samples derived from wildtype *SPB1*, *spb1*$^{D52A}$ and *spb1* $^{D52A/E769K}$ strains.**

| Strain | Cell pellet weight (g) | Final volume (µl) | Final conc. ($OD_{260}$) | Yield (Units/g) |
|---|---|---|---|---|
| *SPB1* prep 1 | 30g | 50 | 8 | 13.3 |
| *SPB1* prep 2 | 30g | 50 | 8 | 13.3 |
| *spb1*$^{D52A}$ prep 1 | 60g | 20 | 3.5 | 1.2 |
| *spb1*$^{D52A}$ prep 2 | 30g | 20 | 2 | 1.3 |
| *spb1* $^{D52A/E769K}$ | 30g | 45 | 8 | 12.0 |

**Supplemental Table S3 - Yeast strains used in this study**

| Strain | Relevant Genotype | Source |
|---|---|---|
| BY4741 | *MATa; his3Δ1; leu2Δ0; met15Δ0; ura3Δ0* | 40 |
| YJE608 | *MATa; his3Δ1; leu2Δ0; met15Δ0; ura3Δ0; trp1Δ::LexA-ED-AD::* TRP1; ura3Δ0::$P_{minCYC1}$-NOG2-Myc-TEV-2xStrep::*URA3; TIF6-3xFLAG-3C-2xProtA::Hph* | this study |
| YJE735 | *MATa; his3Δ1; leu2Δ0; met15Δ0; ura3Δ0; trp1Δ::LexA-ED-AD::* TRP1; ura3Δ0::$P_{minCYC1}$-NOG2-Myc-TEV-2xStrep::*URA3; TIF6-3xFLAG-3C-2xProtA::Hph; spb1D52A::TEF-NrsR-T9* | this study |
| YJE743 | *MATa his3Δ1 leu2Δ0 met15Δ0 ura3Δ0 trp1Δ::LexA-ED-AD::TRP1 SPB1::NrsR TEF-NrsR-T9* | this study |
| YJE744 | *MATa his3Δ1 leu2Δ0 met15Δ0 ura3Δ0 trp1Δ::LexA-ED-AD::TRP1 spb1D52A::NrsR TEF-NrsR-T9* | this study |
| YJE869 | *MATa his3Δ1 leu2Δ0 met15Δ0 ura3Δ0 trp1Δ::LexA-ED-AD::TRP1 spb1D52AE769K::NrsR TEF-NrsR-T9* | this study |
| YJE923 | *MATa; his3Δ1; leu2Δ0; met15Δ0; ura3Δ0; trp1Δ::LexA-ED-AD::* TRP1; ura3Δ0::$P_{minCYC1}$-NOG2-Myc-TEV-2xStrep::*URA3; TIF6-3xFLAG-3C-2xProtA::HYG; spb1D52AE769K::TEF-NrsR-T9* | this study |

### Supplemental Table S4 - Plasmids used in this study

| Plasmid name | Description | Source |
|---|---|---|
| pRS246-GAL | Yeast 2 μ plasmid containing the GAL1 promoter (URA3) | Addgene |
| pJE649 | pFA6a-3xFLAG-3C-6GLY-2xPrtA-HygMX | This paper |
| pJE1047 | pRS426-GAL-*NMD3* | This paper |
| pJE1049 | pRS426-GAL-*NOG2* | This paper |
| pJE1051 | pRS426-GAL-*nog2*$^{G369A}$ | This paper |
| pJE1069 | pRS426-GAL-*SPB1-MTD-NoLS* | This paper |
| pJE1071 | pRS426-GAL-*spb1*$^{D52A}$-*MTD-NoLS* | This paper |

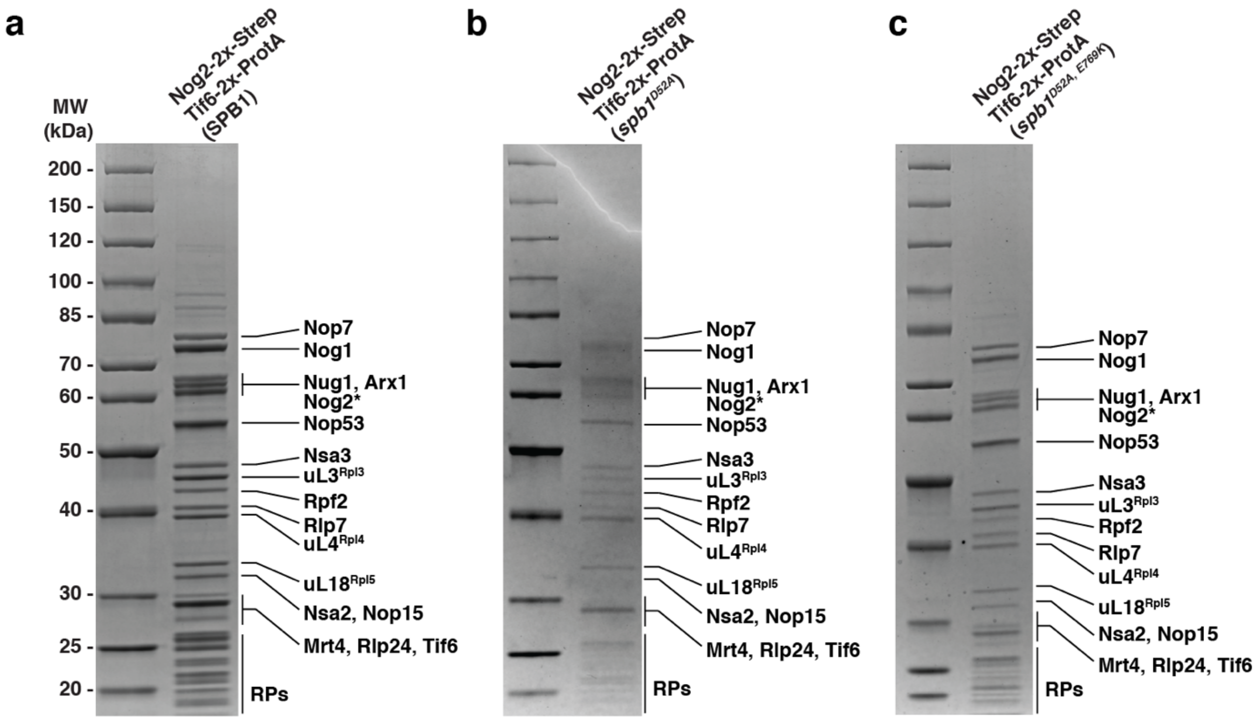


**Supplemental Figure S1 - Purified Nog2·Tif6 pre-ribosomes used for cryo-EM analysis.** **a**, 4 – 20% SDS-PAGE Gel of the sample purified from the wildtype *SPB1* strain. **b**, gel of sample derived from the *spb1*$^{D52A}$ strain. **c**, gel of sample derived from the *spb1*$^{D52A/E769K}$ strain. Protein identification was based on molecular weight, ambiguous bands were identified using mass spectrometry.

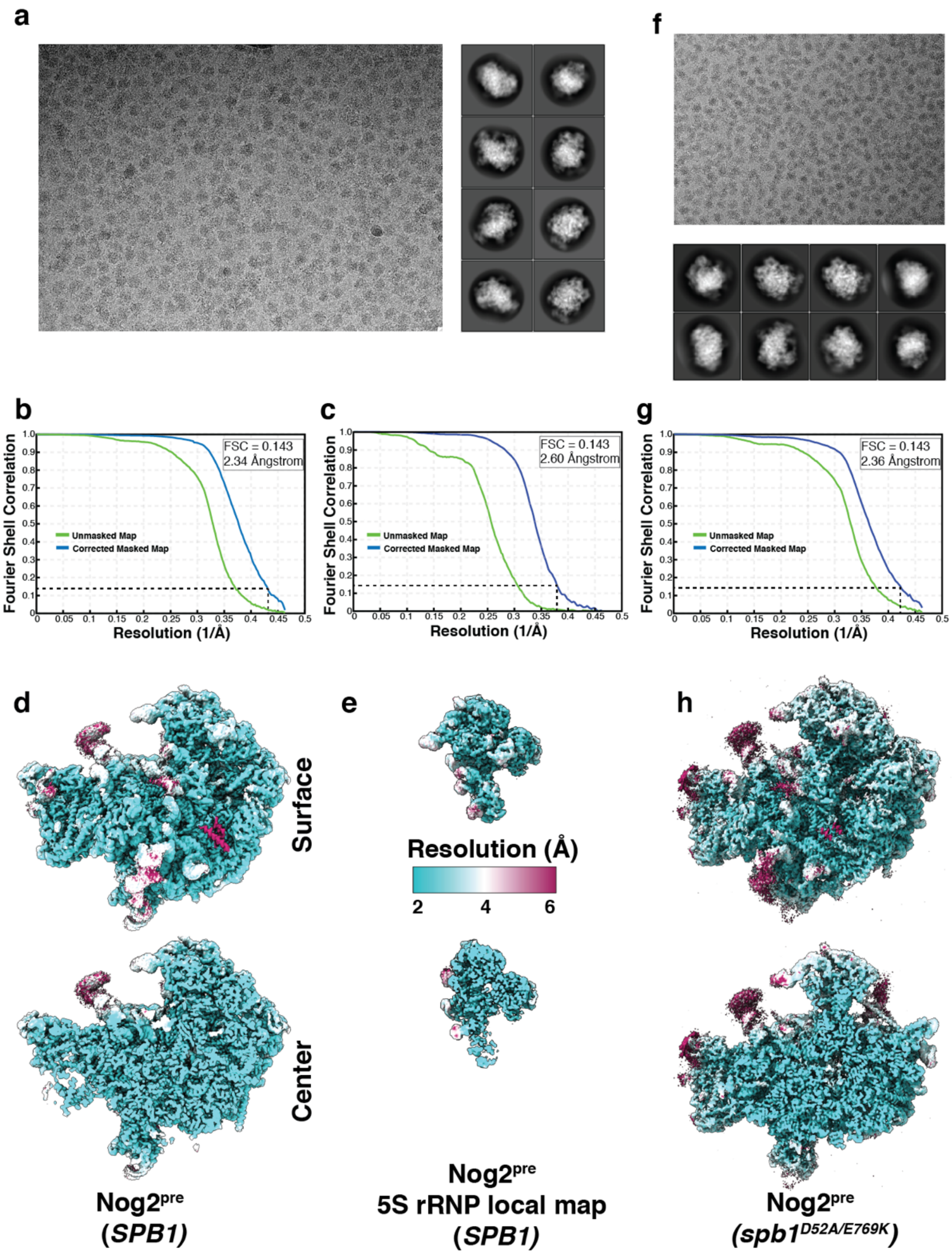


**Supplemental Figure S2 - Micrographs, 2D classes, Fourier shell correlation (FSC) curves and ResMap plots of maps from *SPB1* (panels a-e) or *spb1$^{D52A/E769K}$* (panels f-h) strains.** **a**, representative micrograph and 2D classes. **b**, Fourier shell correlation of unmasked and masked maps with an overall resolution of 2.34 Å. **c**, Fourier shell correlation of unmasked and masked 5S rRNP local map with an overall resolution of 2.60 Å. **d**, overall map (surface and sliced) colored according to local resolution estimates. **e**, resolution distribution of local 5S RNP map. **f**, representative micrograph and 2D classes. **g**, Fourier shell correlation of unmasked and masked maps with an overall resolution of 2.36 Å. **h**, overall pre-rotation map colored according to local resolution estimates.

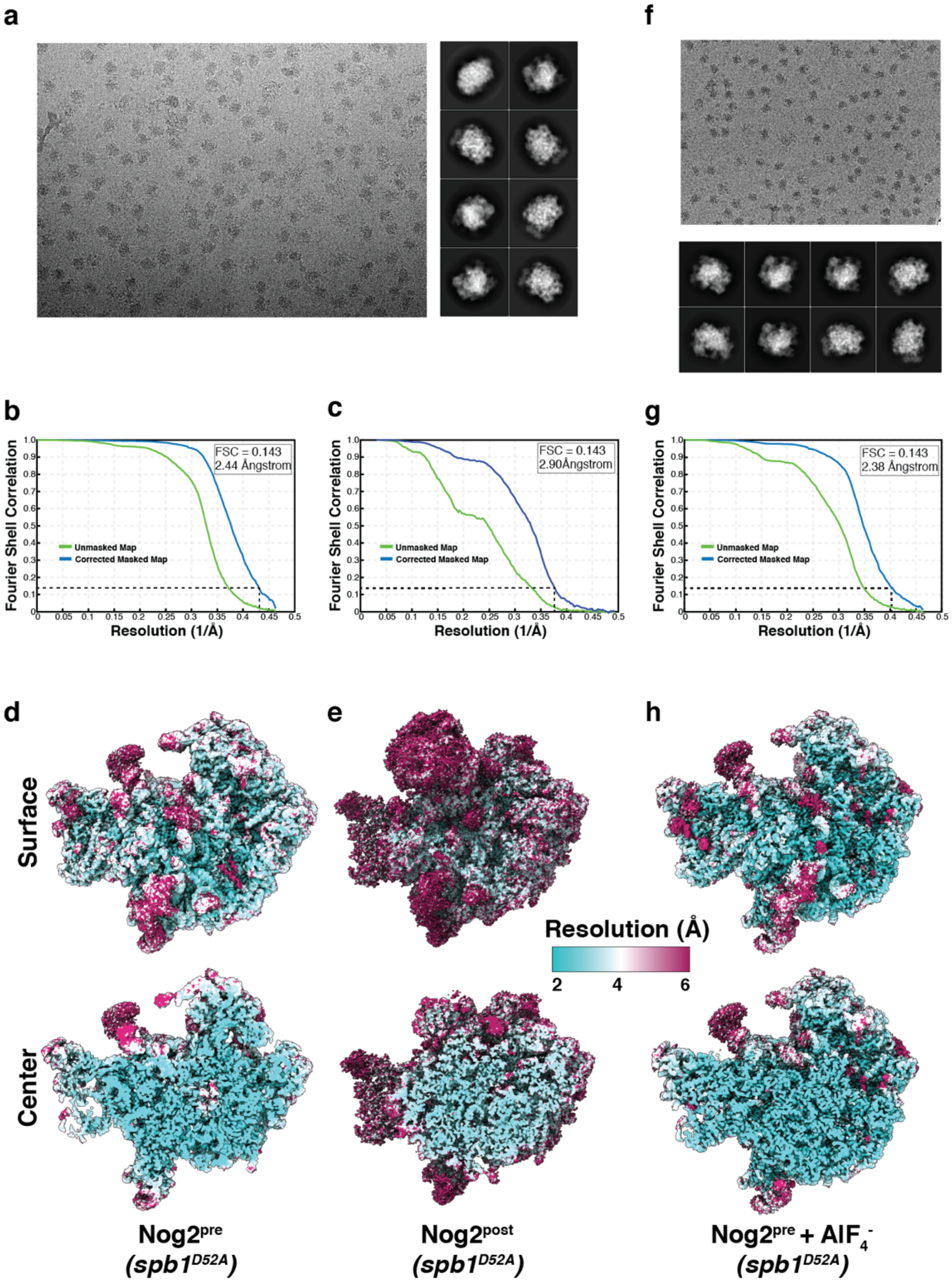


**Supplemental Figure S3 - Micrographs, 2D classes, Fourier shell correlation (FSC) curves and ResMap plots of maps of Nog2$^{pre}$ and Nog2$^{post}$ (panels a-e) and Nog2$^{pre}$+ AlF$_4$- (panels f-h) purified from the *spb1$^{D52A}$* strain**. **a**, representative micrograph and 2D classes. **b**, Fourier shell correlation of unmasked and masked Nog2$^{pre}$ map with an overall resolution of 2.44 Å. **c**, Fourier shell correlation of unmasked and masked Nog2$^{post}$ map with an overall resolution of 2.90 Å. **d**, overall Nog2$^{pre}$ map colored (surface and slice) according to local resolution estimates. **e**, overall Nog2$^{post}$ map is shown and colored according to local resolution estimates. **f**, representative micrograph and 2D classes. **g**, Fourier shell correlation of unmasked and masked Nog2$^{pre}$ map with an overall resolution of 2.38 Å. **h**, overall pre-rotation map colored according to the local resolution estimates.

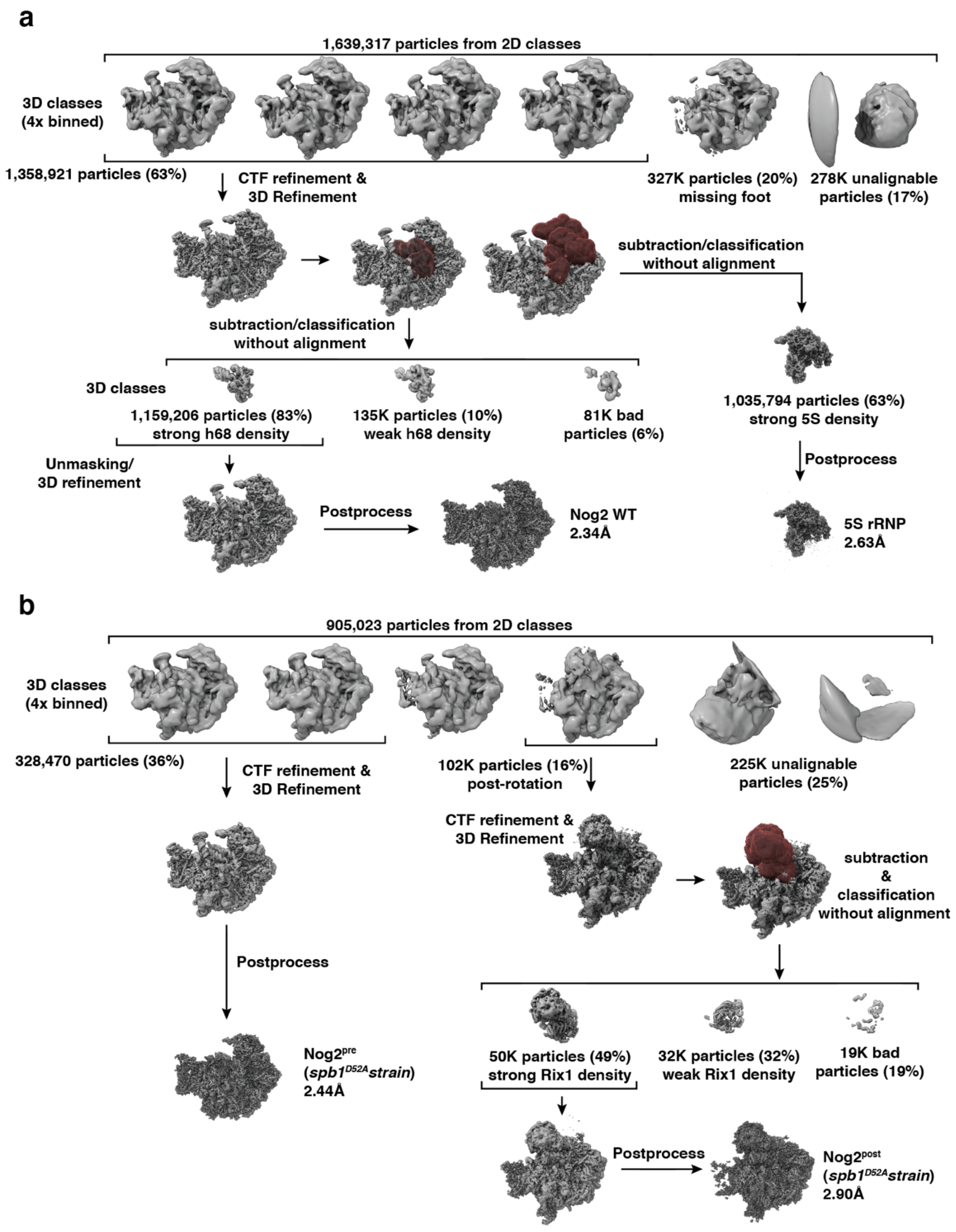


**Supplemental Figure S4 - Cryo-EM data processing, 3D-classification and particle sorting scheme. a**, sorting and refinement scheme for Nog2$^{pre}$ of the wild type *SPB1* sample. **b**, sorting and refinement scheme for Nog2$^{pre}$ and Nog2$^{post}$ of the *spb1$^{D52A}$* sample. Map volumes are shown in gray, masks (dark red) used for subtraction and local classifications are shown aligned to their respective maps. Sorting and classification criteria are denoted, particle number and percentage of the total and final resolution are shown for each map.

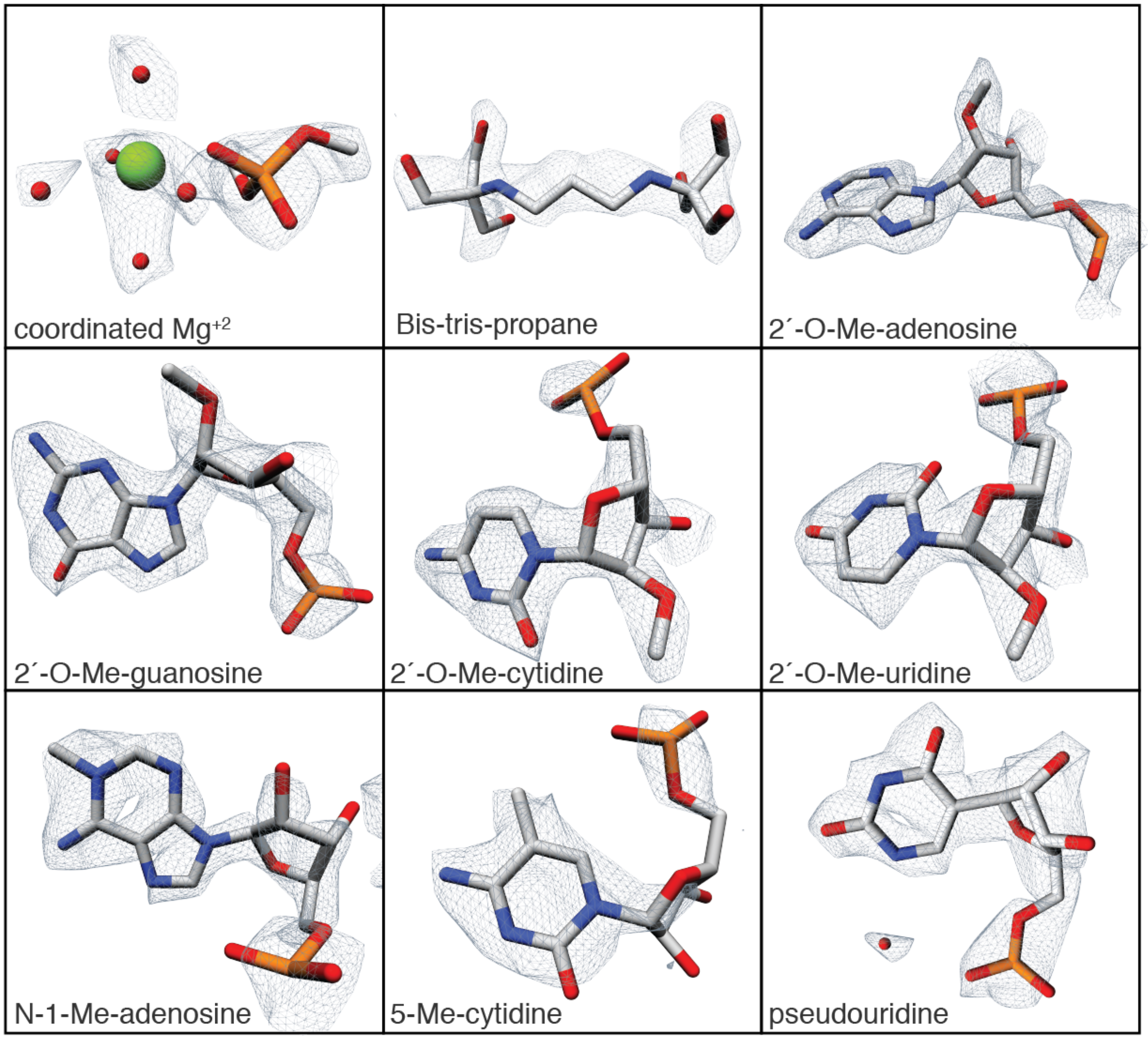


**Supplemental Figure S5 - Representative Cryo-EM map densities for the wild-type Nog2$^{pre}$ sample.** Models and surrounding map density of coordinated magnesium ions, bis-tris-propane molecules and modified nucleotides identified within the Nog2$^{pre}$ maps.

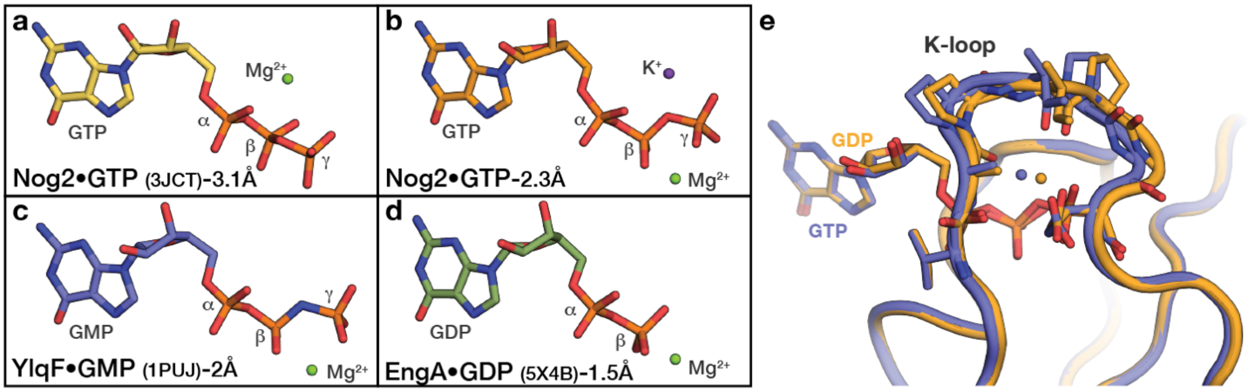


**Supplemental Figure S6 - Map interpretation. a-d**, stick representations of nucleotides in selected K-loop GTPases. **a**, in the original Nog2$^{pre}$ structural model, the γ-phosphate was placed into the density of the $Mg^{2+}$ and the $Mg^{2+}$ ion was placed in the density of the $K^{+}$ ion. **b**, our interpretation of the nucleotide density of Nog2 is consistent with high-resolution crystal structures of bacterial K-loop ATPases YlqF/RbgA and EngA (**c**, **d**). **e**, superposition of Nog2-GTP (blue) and Nog2-GDP (yellow) models shows that GTP hydrolysis causes a ~1Å shift in the position of the K-loop.

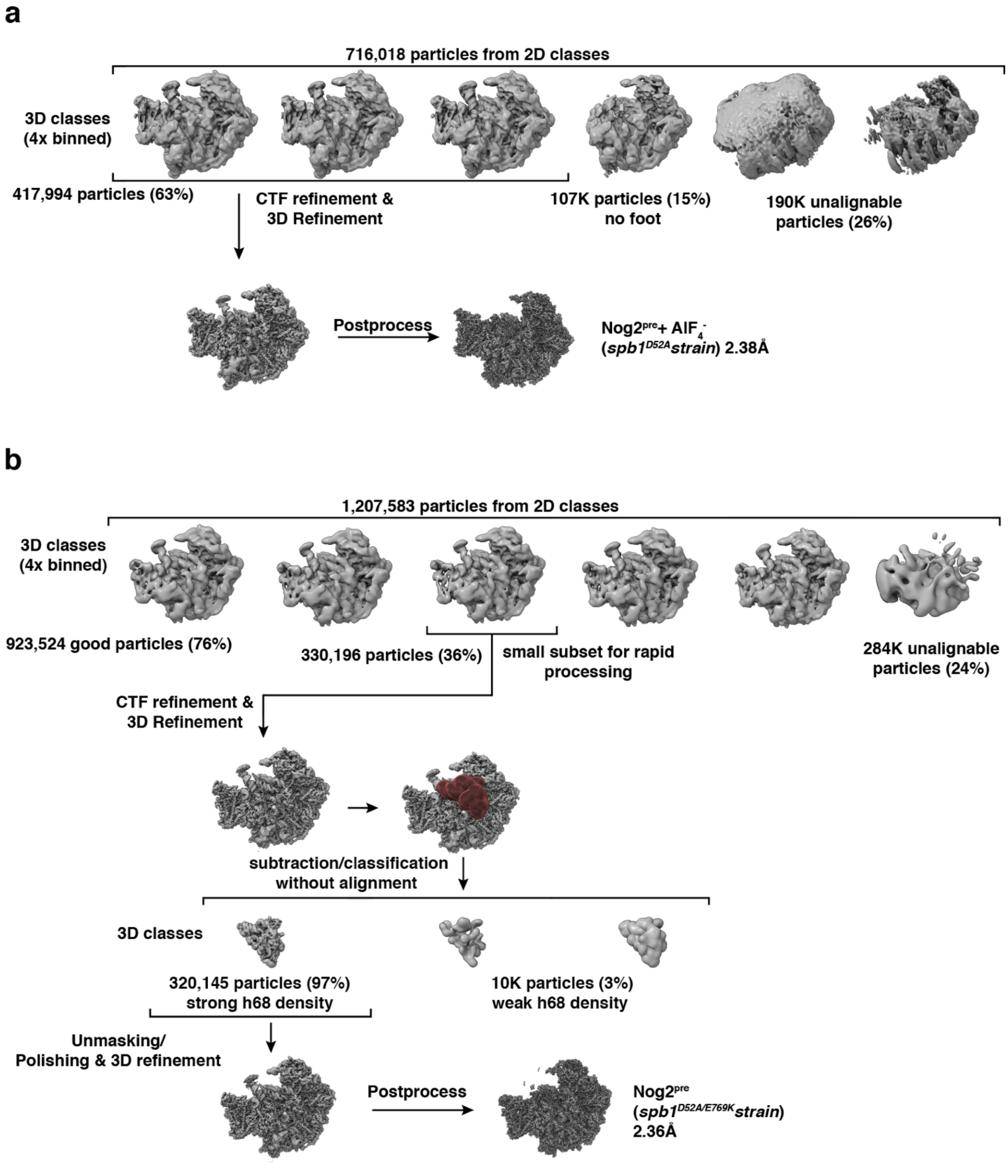


**Supplemental Figure S7 - Cryo-EM data processing, 3D-classification and particle sorting scheme for Nog2·Tif6 intermediates. a**, sorting and refinement scheme for Nog2$^{pre}$ purified from an *spb1*$^{D52A}$ strain in which $AlF_4$- was added before grid preparation. **b**, sorting and refinement scheme for Nog2$^{pre}$ purified from an *spb1*$^{D52A/E769K}$ suppressor strain. Map volumes are shown in gray, masks (dark red) used for subtraction and local classifications are shown aligned to their respective maps. Sorting and classification criteria are denoted, particle number and percentage of the total and final resolution are shown for each map.

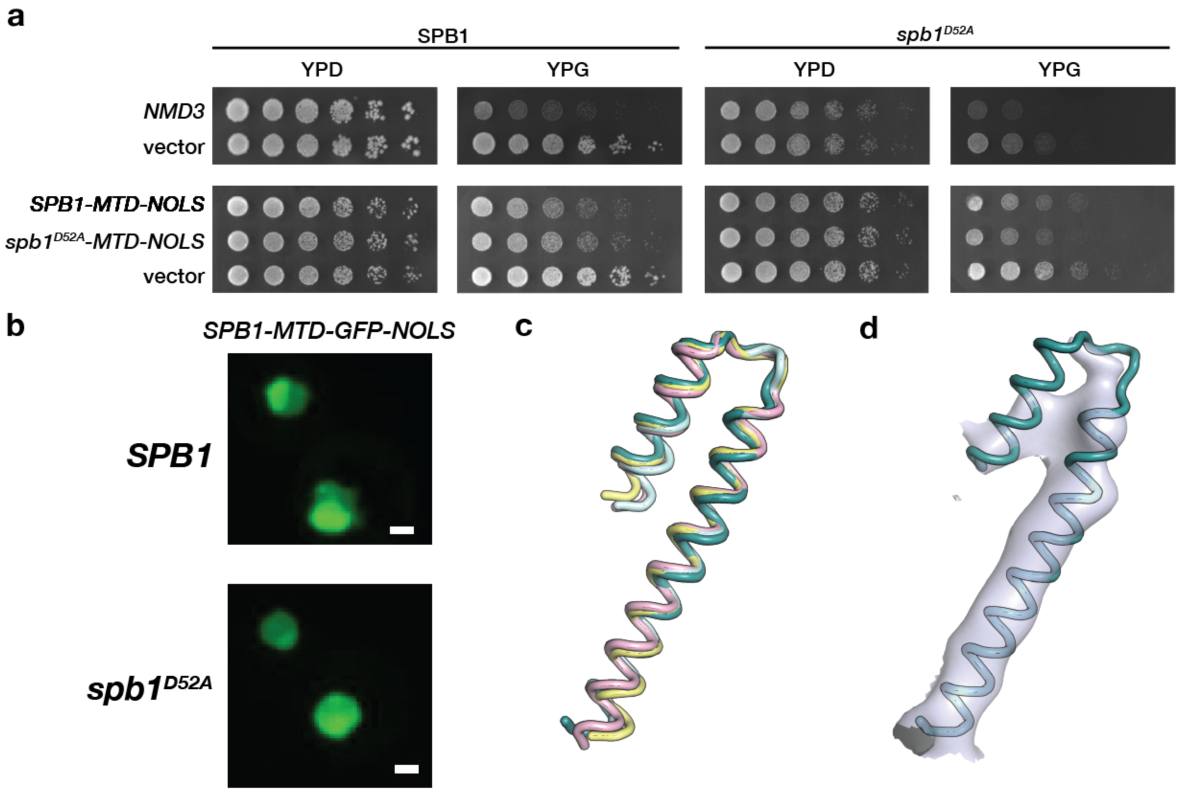


**Supplemental Figure S8 - Overexpression of *NMD3* or *SPB1-MTD* does not rescue growth of *spb1*$^{D52A}$ and location of *spb1*$^{D52A}$ suppressor mutants in the C-terminal domain of Spb1. a,** overexpression of *NMD3* or *SPB1-MTD* does not rescue the slow growth phenotype of the *spb1*$^{D52A}$ strain, instead showing a mild dominant negative effect. Plasmids containing Gal-inducible *NMD3, SPB1-MTD*, *spb1*$^{D52A}$*-MTD* or empty vector were transformed into the *SPB1* or *spb1*$^{D52A}$ strains and plated on Glucose (YPD) or Galactose (YPG). **b**, light microscopy of yeast cells shows the proper nucleolar localization of overexpressed *SPB1-MTD-NOLS*. **c**, superposition of Alphafold models from *S.cerevisiae* (teal), *S.pombe* (pink), *D.melanogaster* (yellow) and *H.sapiens* (cyan) for the helical segment of Spb1 containing the suppressor mutations. **d**, *S.cerevisiae* model docked into the cryo-EM density from the NE1 pre-60S intermediate reconstruction[11].
